# *In vivo* xenogenic reconstitution of human alveolar epithelial architecture and function

**DOI:** 10.64898/2026.01.11.698914

**Authors:** Akira Yamagata, Satoshi Konishi, Satoshi Ikeo, Hiroshi Moriyama, Senye Takahashi, Naoyuki Sone, Satoshi Hamada, Atsushi Saito, Takashi Kawaguchi, Shu Hisata, Akira Niwa, Toshiaki Kikuchi, Hirofumi Chiba, Megumu K. Saito, Koichi Hagiwara, Toyohiro Hirai, Mio Iwasaki, Takuya Yamamoto, Takeshi Takahashi, Shimpei Gotoh

## Abstract

An urgent need exists for lung models that accurately replicate human physiological profiles. We developed a chimeric mouse model enabling targeted ablation of alveolar type 2 (AT2) cells and lung macrophages, creating niches for endoscopically transplanted human induced pluripotent stem cell (hiPSC)-derived lung progenitors (hLPs). These engrafted cells were retained for 24 weeks, demonstrating self-renewal and differentiation potential, surfactant protein secretion, and maintaining alveolar phosphate homeostasis, suggesting their maturation into AT2 cells. Furthermore, transplantation of hLPs derived from disease-specific hiPSCs recapitulated the phenotype of pulmonary alveolar microlithiasis. The architecture, function, and metabolism of human alveolar epithelium were accurately replicated *in vivo.* This model has the potential to link experimental models and first-in-human studies, facilitating the development of novel therapies for intractable lung diseases.

## Main Text

Chronic lung diseases like chronic obstructive pulmonary disease and interstitial pneumonia, as well as acute respiratory diseases such as bacterial pneumonia and COVID-19, are significant global causes of mortality. The limited regenerative capacity of lungs and the lack of effective treatments highlight the urgent need for new therapies, including regenerative medicine. However, the translation of preclinical findings into clinical practice is hindered by the absence of accurate *in vivo* models of the human lung environment. Significant differences in genome, cell types, and lung anatomy between humans and mice limit the effectiveness of mouse models for simulating human diseases (*1*, *2*).

Advances in three-dimensional organoid culture techniques have enabled the *in vitro* expansion of alveolar epithelial cells using human primary cells (*3*–*8*) or pluripotent stem cells (*9*–*11*). Particularly, human induced pluripotent stem cells (hiPSCs) are used to model inheritable lung diseases involving alveolar epithelial cell disorders (*12*–*16*). However, replicating the lungs’ complex, three-dimensional structure, comprising various cell types, including epithelial, mesenchymal, vascular endothelial, and immune cells, interacting within an organ system, and influenced by physiological factors via the blood, lymphatic, and nervous systems, is challenging.

To overcome these challenges, researchers have developed humanized animal models by introducing specific human cells and tissues into immunocompromised animals to replicate human physiological and metabolic profiles. Numerous studies have documented the transplantation of human epithelial cells into the injured lungs of immunocompromised mice to reconstruct airway (*17*, *18*) and alveolar (*19*–*21*) epithelial cells. We previously reported that xenotransplantation of human lung progenitor cells (hLPs) derived from hiPSCs into the left lungs of immunocompromised mice results in their differentiation into alveolar and airway epithelial cells (*22*). However, due to the limited replacement rate of endogenous epithelial cells and short evaluation periods, the physiological function of reconstituted alveolar epithelial cells and their application to disease modeling have not been extensively investigated.

In the current study, we developed an efficient method for transplanting hiPSC-derived hLPs and alveolar stem cells using lung-humanized mice to analyze alveolar epithelial cell function and model lung diseases. This platform can be used in future studies to define human lung cell functions and aid in developing new lung disease therapies.

### Durable orthotopic engraftment of hiPSC-derived human lung progenitor cells in *NOG-LysM-DTR* mice

Administering diphtheria toxin (DT) to *LysM-DTR* mice—expressing the human DT receptor (DTR) under *lysozyme M* (*LysM*) promoter control—efficiently ablates lung-resident macrophages and alveolar type 2 (AT2) cells (*23*, *24*), i.e., tissue stem cells in alveoli (*25*). To create a niche for the post-transplant proliferation and differentiation of human alveolar epithelial cells in immunocompromised mice, *NOD/Shi-scid, IL-2Rγ KO Jic* (*NOG*) mice were crossed with *LysM-DTR* mice to generate *NOG-LysM-DTR* mice.

DT was injected at various doses into the left main bronchus of *NOG-LysM-DTR* mice using an endoscope-assisted transtracheal administration system (EATAS) (Fig. 1A, fig. S1, A and B, and Movie S1), as previously reported (*22*). DT caused weight loss and death in mice in a dose-dependent manner (fig. S1, C to E). In contrast, no mice died in the vehicle control group. Macroscopic observation revealed hemorrhaging in the left lobe, while the right lobes remained relatively intact at three weeks post-DT administration (Fig. 1B). Additionally, four days after DT administration, the number of AT2 cells (proSPC^+^ NaPi2b^+^) and macrophages (CD68^+^) decreased in a dose-dependent manner in the left lobe, while maintaining alveolar structure without marked morphological change (Fig. 1, C to E, and fig. S1, F and J). Similar effects were not observed in the right lobes. Although significant tissue damage and fibrosis were observed in DT-treated left lobe three weeks post-administration (fig. S1, F to H), proSPC and NaPi2b expression were restored (fig. S1I). This suggested that the left lobe was undergoing regeneration after injury. Collectively, these results highlighted the selective ablation of AT2 cells and macrophages in the left lobe of *NOG-LysM-DTR* mice.

**Fig. 1.**
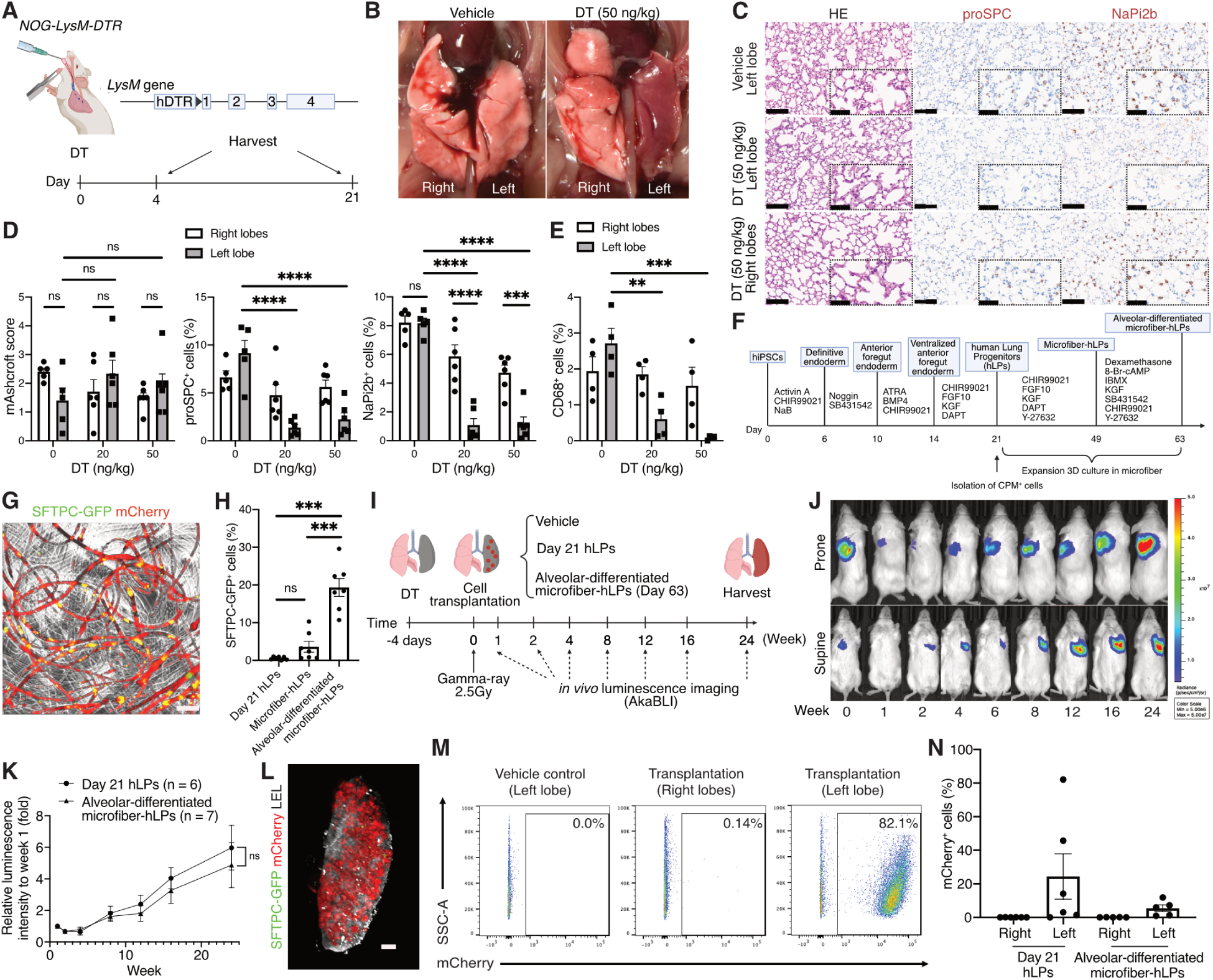
Long-term orthotopic engraftment of hiPSC-derived lung epithelial cells in *NOG-LysM-DTR* mice. (A) Schematic of diphtheria toxin (DT) administration into the left main bronchus of *NOG-LysM-DTR* mice and lung harvesting. (B) Macroscopic observation of *NOG-LysM-DTR* mice 3 weeks after DT or vehicle administration into the left main bronchus. (C) Hematoxylin and eosin (H&E) staining and immunohistochemistry (IHC) of lung sections of mice 4 days after DT treatment. Scale bars, 100 μm and 50 μm. (D) Assessment of the modified Ashcroft score, proSPC^+^ cells and NaPi2b^+^ cells 4 days after DT treatment (mean ± SEM) (n = 5–6/group). Statistical comparisons were performed using two-way repeated measures ANOVA with Turkey’s post-hoc analysis. ns: not significant, **P < 0.01, ***P < 0.001, ****P < 0.0001. (E) Assessment of CD68^+^ cells 4 days after DT treatment (mean ± standard error of the mean [SEM]); n = 4/group). Statistical comparisons were performed using two-way repeated measures ANOVA with Turkey’s post-hoc analysis. **P < 0.01, ***P < 0.001. (F) Schematic diagram of the stepwise differentiation and expansion from hiPSCs into alveolar-differentiated microfiber-hLPs. (G) Live imaging of alveolar-differentiated microfiber-hLPs. Scale bar, 1 mm. (H) Flow cytometric quantification of SFTPC-GFP^+^ cell proportions in day 21 hLPs, microfiber-hLPs, and alveolar-differentiated microfiber-hLPs (mean ± SEM; n = 7 from seven independent experiments). Statistical comparisons were performed using one-way repeated measures ANOVA with Turkey’s post-hoc analysis. ns: not significant, ***P < 0.001. (I) Experimental strategy for transplantation. (J) Representative serial pictures of *in vivo* luminescence imaging of a mouse transplanted with day 21 hLPs following injury at DT 20 ng/kg. (K) Relative intensity of bioluminescence for 24 weeks after transplantation of day 21 hLPs (n = 6) and alveolar-differentiated microfiber-hLPs (n = 7) following injury at DT 20 ng/kg, with the luminescence intensity at week one as 1 (mean ± SEM). Statistical comparisons were performed using two-way repeated measures ANOVA with Sidak’s post-hoc analysis. ns: not significant. (L) 3D image of cleared left lung 24 weeks after transplantation of day 21 hLPs rendered on Imaris. Scale bar, 1 mm. See Movie S2. (M and N) Flow cytometric quantification of the proportion of mCherry^+^ cells among CD31^-^CD45^-^ cells in murine lungs 24 weeks post-transplantation of day 21 hLPs and alveolar-differentiated microfiber-hLPs (mean ± SEM) (n = 5–6), respectively.

Traditionally, assessing transplanted cell viability in deep organs like the lungs has required post-mortem analysis. AkaBLI, a new luminescence system, enables *in vivo* single-cell imaging in deep organs (*26*). To monitor transplanted cells noninvasively, an *SFTPC^GFP^* knock-in reporter hiPSC line (B2-3) (*27*) was modified to stably express *mCherry-Akaluc* by inserting the cassette into the *AAVS1* safe-harbor locus using CRISPR-Cas9-mediated gene editing (maB2-3) (fig. S1K). The maB2-3 clone maintained a normal karyotype (fig. S1L). Subsequently, maB2-3 was differentiated into ventralized anterior foregut endodermal cells and NKX2.1^+^ hLPs were isolated using an anti-Carboxypeptidase M (CPM) antibody on day 21 (day 21 hLPs) (*9*, *27*) (Fig. 1F). For efficient transplantation, day 21 hLPs were cultured on non-adherent plates for an additional two days to form aggregates (fig. S1M).

To evaluate the benefit of transplanting differentiated alveolar lineage cells, day 21 hLPs were expanded for four weeks using a cell-laden core-shell microfiber system (microfiber-hLPs) and differentiated into alveolar epithelial cells over two additional weeks (alveolar-differentiated microfiber-hLPs), as previously reported (*22*). During differentiation and expansion, the proportion of SFTPC-GFP^+^ cells among the total mCherry^+^ cells increased (Fig. 1, G and H, and fig. S1, N and O), confirming their maturation into alveolar lineages.

Single-cell RNA sequencing was performed of FACS-sorted mCherry^+^ cells in alveolar-differentiated microfiber-hLPs to further evaluate the cell population (fig. S1, P to R). The resulting dataset was visualized with UMAP. Clusters were annotated based on lung cell-type gene signatures. Some cell clusters expressed the gene signatures of alveolar epithelial cells and accounted for approximately 50% of all cells. Among them, a certain cluster expressed proliferation markers such as *TOP2A* and *MKI67*.

Four days after DT treatment (20 or 50 ng/kg), *NOG-LysM-DTR* mice were irradiated with 2.5 Gy γ-ray to eliminate proliferating endogenous cells. Subsequently, day 21 hLPs or alveolar-differentiated microfiber-hLPs were transplanted into the left main bronchus of *NOG-LysM-DTR* mice using EATAS. (Fig. 1I). The transplanted cells emitted luminescence from the left lung for at least 24 weeks (Fig. 1, J and K, and fig. S2A). The luminescence intensity from the engrafted cells decreased for the first 2 weeks but then increased, suggesting *in vivo* proliferation of the engrafted cells. The luminescence intensity of transplanted alveolar-differentiated microfiber-hLPs at 20 ng/kg DT did not differ significantly from that at 50 ng/kg DT (fig. S2B). Three days post-transplantation of alveolar-differentiated microfiber-hLPs, small, scattered clusters of mCherry^+^ cells appeared, growing into larger clusters by 105 days (fig. S2C).

After clearing the resected lungs 24 weeks post-transplantation of day 21 hLPs using the CUBIC method (*28*), 3D imaging revealed extensive mCherry^+^ human cell engraftment in the left lobe only (Fig. 1L, fig. S2D, and Movie S2). Tissue sections confirmed that these alveolar-differentiated microfiber-hLPs settled within the left lobe without impacting the alveolar structure (fig. S2E). Flow cytometry revealed that mCherry^+^ cells comprised 24.40 ± 13.50% (mean ± SEM) of day 21 hLPs and 5.46 ± 2.03% (mean ± SEM) of alveolar-differentiated microfiber-hLPs among the CD31^-^CD45^-^ cells in the left lung (Fig. 1, M and N, and fig. S2F). Thus, engrafted human cells were structurally integrated with the host lung.

### Transplanted cells differentiate into alveolar epithelial cells *in vivo*

To evaluate the transcriptional profiles of engrafted cells, single-cell RNA sequencing was performed of FACS-sorted mCherry^+^ cells at 24 weeks post-transplantation of day 21 hLPs or alveolar-differentiated microfiber-hLPs (Fig. 2A). mCherry^-^ cells were excluded as endogenous cells. To prevent contamination from mouse transcripts, reads were mapped hg38 and mm39; only those uniquely aligned to hg38 were analyzed. The resulting datasets were visualized with UMAP (fig. S3A). All clusters were verified to express *mCherry-Akaluc* (fig. S3B).

**Fig. 2.**
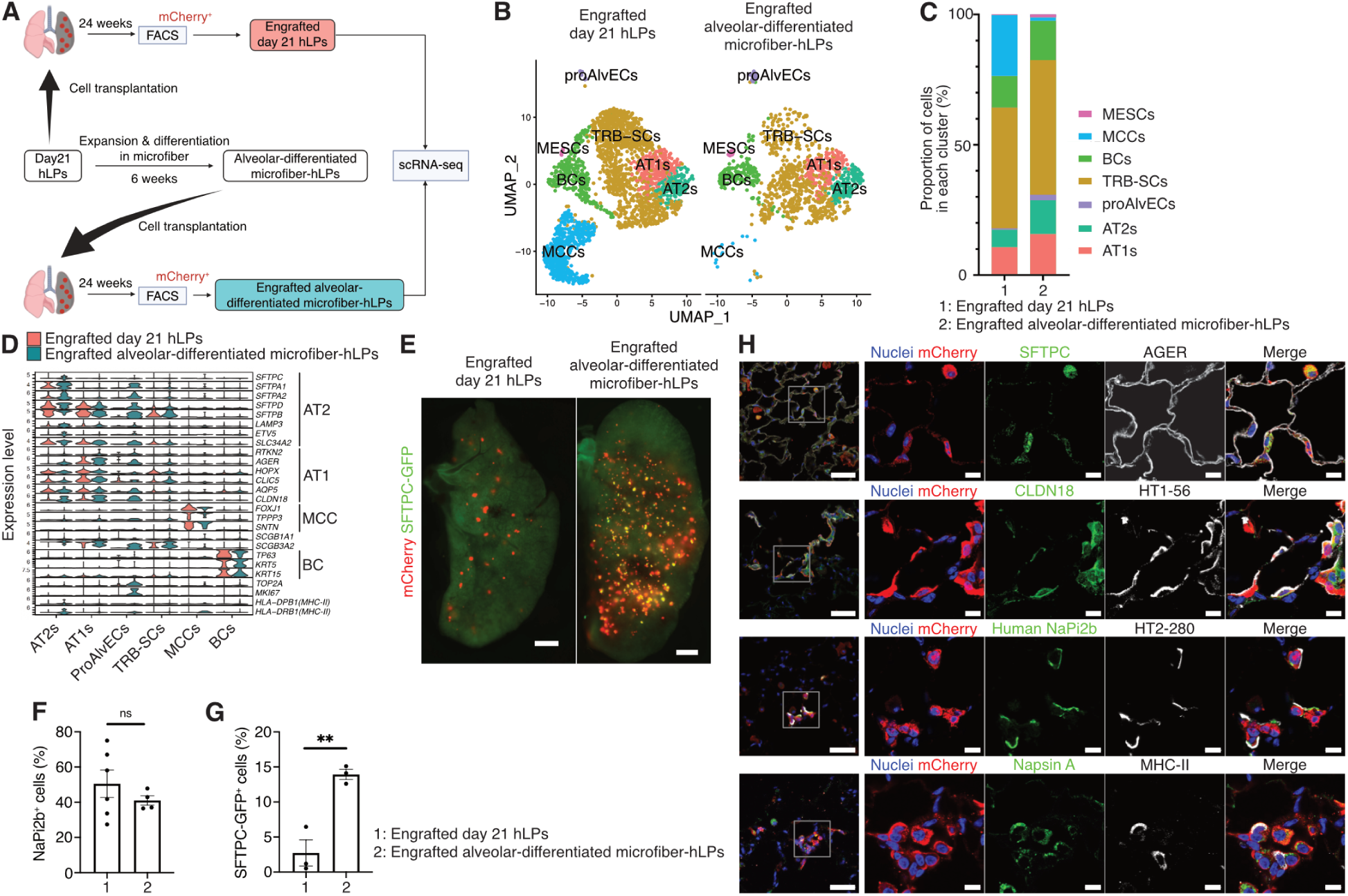
Transplanted hLPs differentiate into alveolar epithelial cells *in vivo*. (A) Experimental strategy for scRNA-seq analysis of engrafted day 21 hLPs and alveolar-differentiated microfiber-hLPs 24 weeks post-transplantation. (B) UMAP plot of integrated scRNA-seq data from engrafted cells derived from day 21 hLPs (left) and alveolar-differentiated microfiber-hLPs (right), visualized using split view. Abbreviations used in the figure: proAlvECs, proliferative alveolar epithelial cells; TRB-SCs, terminal and respiratory bronchiole-secretory cells; BCs, basal cells; MESCs, mesenchymal cells. (C) Bar plot showing the proportion of cells in each cluster for two transplantation conditions: engrafted day 21 hLPs (1-left) and engrafted alveolar-differentiated microfiber-hLPs (2-right). (D) Stacked violin plots of selected marker gene expression across clusters, split by condition: engrafted day 21 hLPs (left) and alveolar-differentiated microfiber-hLPs (right). (E) Representative live imaging of mouse left lungs 3 weeks post-transplantation of day 21 hLPs and alveolar-differentiated microfiber-hLPs, respectively. Scale bar, 1 mm. (F) Flowcytometric quantification of NaPi2b^+^ cell proportions among mCherry^+^ cells in murine left lungs 24 weeks post-transplantation of day 21 hLPs or alveolar-differentiated microfiber-hLPs (mean ± SEM) (n = 4–6); unpaired two-tailed t-test, ns: not significant. (G) Flowcytometric quantification of SFTPC-GFP^+^ cell proportion among mCherry^+^ cells in murine left lungs 24 weeks post-transplantation of day 21 hLPs or alveolar-differentiated microfiber-hLPs (mean ± SEM) (n = 3); unpaired, two-tailed t-test, **P < 0.01. (H) Immunofluorescence imaging of *NOG-LysM-DTR* mouse left lung after transplanting alveolar-differentiated microfiber-hLPs. The leftmost images show low-power field views. Scale bars, 50 μm and 10 μm, magnified views.

All human cells from each dataset were combined and plotted with UMAP (fig. S3C). The integrated UMAP dataset was divided into cell clusters based on the Louvain algorithm, with similar transcriptional profiles manually merged (Fig. 2B). Certain cell clusters expressed high levels of AT2 or alveolar type 1 (AT1) gene signatures without other cell-type signatures. The proportions of AT2 and AT1 cells were higher in engrafted alveolar-differentiated microfiber-hLPs than in the engrafted day 21 hLPs, whereas multi-ciliated cells (MCCs) comprised a larger proportion of day 21 hLPs (Fig. 2C). Notably, AT2 cells in engrafted alveolar-differentiated microfiber-hLPs expressed *SFTPC* and major histocompatibility complex II *(MHCII)*-related mRNA, which is expressed specifically in AT2 cells (*29*–*32*), not day 21 hLPs (Fig. 2D and fig. S3D). Moreover, AT1 cells in the engrafted alveolar-differentiated microfiber-hLPs exhibited higher expression of AT1 profile genes *RTKN2* and *AGER* than engrafted day 21 hLPs.

To investigate the effect of murine lung microenvironment on the maturation of AT2 cells, we also integrated cell clusters corresponding to AT2 cells (AT2s and imAT2s) from the dataset obtained before transplantation of alveolar-differentiated microfiber-hLPs with those from the two datasets obtained 24 weeks after transplantation and compared the gene expression of AT2 markers (fig. S3E). Engrafted alveolar-differentiated microfiber-hLPs exhibited higher expression of AT2 profile genes (*SFTPA1*, *SFTPA2*, *SFTPB*, *SFTPC*, *LAMP3*, *SLC34A2*, and *HLA-DRB1*) compared to pre-transplant alveolar-differentiated microfiber-hLPs, suggesting that the host microenvironment promotes AT2 maturation. Collectively, these findings suggest that transplanted alveolar-differentiated microfiber-hLPs primarily differentiate into cells with gene expression patterns similar to primary adult human AT2 and AT1 cells.

Three weeks after transplantation, fluorescence microscope live imaging revealed that SFTPC-GFP was expressed by few engrafted day 21 hLPs in the left lobe and many alveolar-differentiated microfiber-hLPs (Fig. 2E). After 24 weeks, flow cytometry detected NaPi2b expression by approximately 40% of engrafted day 21 hLPs and alveolar-differentiated microfiber-hLPs (Fig. 2F and fig. S3F). Meanwhile, over 10% of the engrafted alveolar-differentiated microfiber-hLPs were SFTPC-GFP^+^, significantly more than day 21 hLPs (Fig. 2G and fig. S3F). Hematoxylin and eosin staining further revealed no signs of malignancy 24 weeks post-transplantation (fig. S3G). Additionally, NaPi2b expression was localized toward the alveolar lumen, demonstrating maintained apico–basal polarity of the engrafted cells within the host tissue context.

Immunofluorescence staining of the left lobe following alveolar-differentiated microfiber-hLP transplantation revealed SFTPC, NaPi2b, HT2-280, Napsin A, and MHC-II expression—AT2 cell markers—in mCherry^+^ cells. This suggests that alveolar-differentiated microfiber-hLPs retained AT2 markers expression *in vivo* (Fig. 2H). Additionally, HT1-56 and AGER—AT1 cell markers—were also expressed by elongated forms of mCherry^+^ cells, suggesting their maturation into AT1 cells (Fig. 2H and fig. S3H). Furthermore, mCherry^+^ cells expressed CLDN18—a AT1-specific tight junction protein—suggesting that the engrafted cells reconstructed alveolar barrier integrity *in vivo*.

To investigate the secretory function of the engrafted cells, the bronchoalveolar lavage fluid (BALF) collected 24 weeks post-transplantation from *NOG-LysM-DTR* mice was subjected to proteomic analysis. After excluding proteins with peptides common to humans and mice, 125 human-specific proteins were identified. Human SFTPB was detected in 50% (2/4) mice transplanted with day 21 hLPs, and 80% (4/5) of those transplanted with alveolar-differentiated microfiber-hLPs (fig. S3I). Meanwhile, human SFTPD was absent in the BALF of day 21 hLP-transplanted mice but was detected in 20% (1/5) of those transplanted with alveolar-differentiated microfiber-hLPs (fig. S3J). Moreover, human SCGB1A1 was detected in 75% (3/4) of BALF samples from day 21 hLP-transplanted mice and 100% (5/5) of mice transplanted with alveolar-differentiated microfiber-hLPs (fig. S3K). These findings suggest that engrafted cells secreted lung-specific proteins into the alveolar space *in vivo*.

### Monolayer culture of alveolar-differentiated microfiber-hLPs recapitulates the barrier integrity, cell polarity, and phosphate uptake function of alveolar epithelial cells

AT2 cells produce pulmonary surfactant to normalize surface tension at the air–liquid interface in alveoli. In the alveolar fluid compartment, phosphate, a major surfactant constituent, is efficiently taken up and recycled by AT2 cells via NaPi2b—a major sodium-dependent phosphate cotransporter on the apical membrane (*33*, *34*). AT1 and AT2 cells form the alveolar epithelial barrier via tight junctions and adherens junctions, preventing fluid leakage into the airspace (*35*, *36*). Thus, hLPs derived from hiPSCs were used to recapitulate the barrier integrity and polarity of alveolar epithelial cells and the phosphate uptake function of AT2 cells *in vitro* and *in vivo*.

First, to model the apico–basal polarity, intercellular junctions, and alveolar phosphate metabolism *in vitro*, the alveolar-differentiated microfiber-hLPs were dissociated and seeded onto Geltrex-coated cell culture inserts (Fig. 3A). The effects of this culture on alveolar barrier function were investigated, revealing that transepithelial electrical resistance (TEER) gradually increased by 7 days post-passage (Fig. 3B). Immunofluorescence staining of reseeded alveolar-differentiated microfiber-hLPs derived from *SFTPC^GFP^* hiPSCs after three days of culturing revealed a monolayer structure, with NaPi2b localized to the apical side (Fig. 3C). The tight junction protein zonula occludens-1 (ZO-1) was expressed between the cells (fig. S4A).

**Fig. 3.**
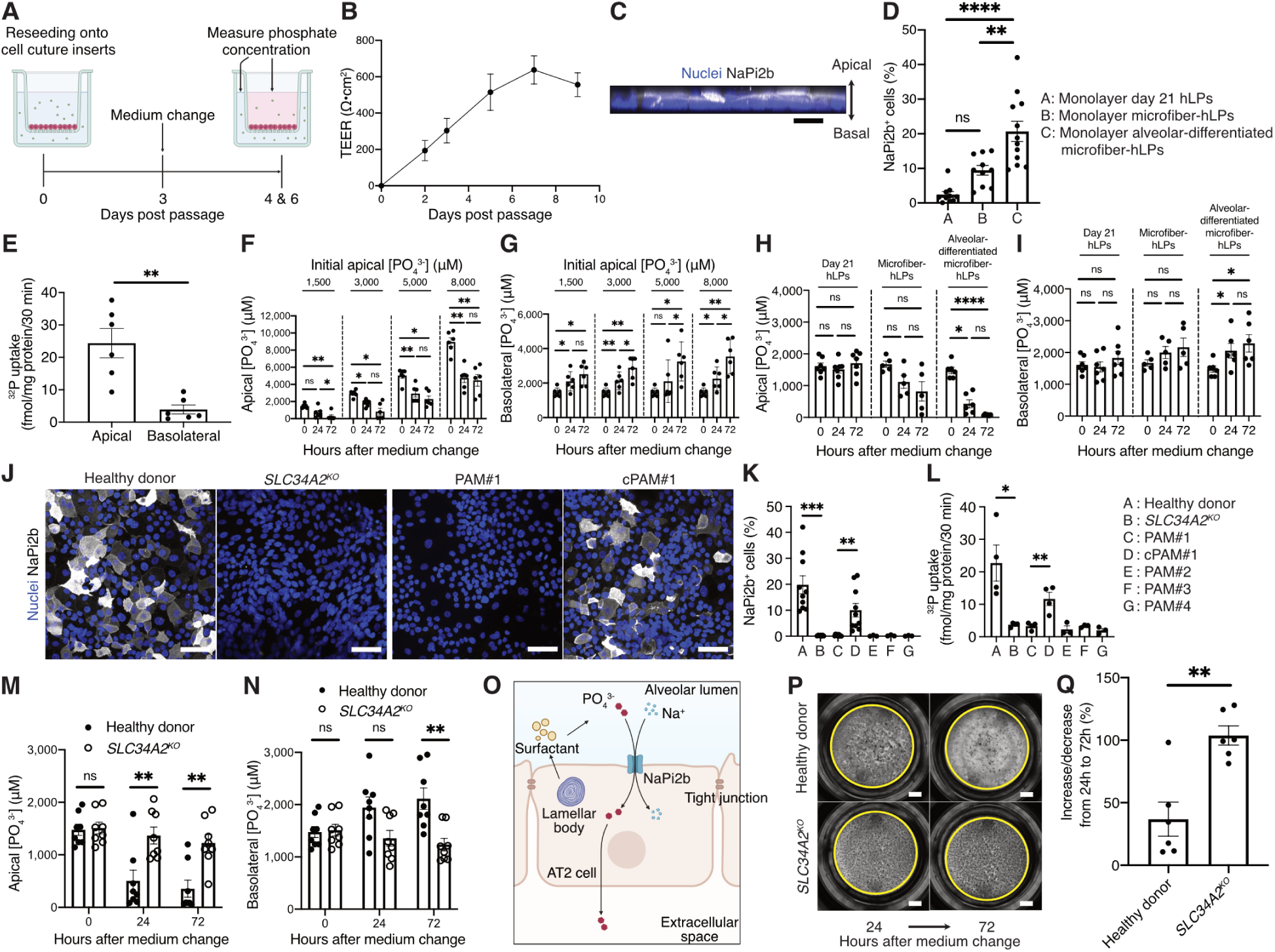
Functional analysis of alveolar-differentiated microfiber-hLPs enables pulmonary alveolar microlithiasis modeling. (A) Experimental strategy for monolayer culture of alveolar-differentiated microfiber-hLPs on cell culture inserts for harvesting apical and basolateral medium. (B) Transepithelial electrical resistance (TEER) in monolayer cultures of reseeded alveolar-differentiated microfiber-hLPs derived from *SFTPC^GFP^*hiPSCs (B2-3) (n = 6). Data are expressed as mean ± standard error of the mean (SEM). (C) Representative immunofluorescence image of the apico-basal axis in the alveolar-differentiated microfiber-hLPs cultured on a cell culture insert for 3 days. Scale bar, 20 µm. (D) NaPi2b^+^ cell proportion in the total day 21 hLPs, microfiber-hLPs (day 49), and alveolar-differentiated microfiber-hLPs (day 63) three days after passaging onto cell culture inserts (n = 10–12/group); one-way analysis of variance (ANOVA) with Turkey’s post-hoc analysis, ns: not significant, **P < 0.01, ****P < 0.0001. (E) ^32^P uptake from apical or basolateral medium into alveolar-differentiated microfiber-hLPs derived from *SFTPC^GFP^* hiPSCs (B2-3) 3 days after passaging onto cell culture inserts. (n = 6/group from six independent experiments). Data are expressed as mean ± SEM; paired, two-tailed t-test, **P < 0.01. (F) Colorimetric assay of phosphate concentrations in apical medium of monolayer culture at 0 h, 24 h, 72 h in alveolar-differentiated microfiber-hLPs derived from *SFTPC^GFP^* hiPSCs after medium change prepared to apical phosphate concentrations of 1500 μM, 3000 μM, 5000 μM, and 8000 μM (n = 6/group). Data are expressed as mean ± SEM; one-way repeated measures ANOVA with Turkey’s post-hoc analysis, ns: not significant, *P < 0.05, **P < 0.01 (G) Colorimetric assay of phosphate concentrations in basolateral medium of monolayer culture at 0 h, 24 h, 72 h in alveolar-differentiated microfiber-hLPs derived from *SFTPC^GFP^* hiPSCs after medium change prepared to apical phosphate concentrations of 1500 μM, 3000 μM, 5000 μM, and 8000 μM (n = 6/group). Data are expressed as mean ± SEM; one-way repeated measures ANOVA with Turkey’s post-hoc analysis, ns: not significant, *P < 0.05, **P < 0.01 (H) Colorimetric assay of phosphate concentrations in apical medium of monolayer culture at 0 h, 24 h, 72 h after medium change in day 21 hLPs, microfiber-hLPs, and alveolar-differentiated microfiber-hLPs derived from *SFTPC^GFP^* hiPSCs (n = 5–7/group). Data are expressed as mean ± SEM; one-way repeated measures ANOVA with Turkey’s post-hoc analysis, ns: not significant, *P < 0.05, ****P < 0.0001. (I) Colorimetric assay of phosphate concentrations in basolateral medium of monolayer cultures at 0 h, 24 h, 72 h after medium change in day 21 hLPs, microfiber-hLPs and alveolar-differentiated microfiber-hLPs derived from *SFTPC^GFP^*hiPSCs (n = 5–7/group); mean ± SEM; one-way repeated measures ANOVA with Turkey’s post-hoc analysis, ns: not significant, *P < 0.05. (J) Immunofluorescence images in alveolar-differentiated microfiber-hLPs derived from *SFTPC^GFP^* hiPSCs, *SLC34A2^KO^* hiPSCs, PAM#1-hiPSCs, and cPAM#1-hiPSCs reseeded and cultured on cell culture inserts for 3 days. Scale bar, 50 µm. (K) Proportion of NaPi2b^+^ cells in total alveolar-differentiated microfiber-hLPs derived from *SFTPC^GFP^* hiPSCs, *SLC34A2^KO^* hiPSCs, PAM#1-4-hiPSCs, and cPAM#1-hiPSCs three days after passaging onto cell culture inserts (n = 3–10/group). Data are expressed as mean ± SEM; unpaired, two-tailed t-test, **P < 0.01, ***P < 0.001. (L) ^32^P uptake into alveolar-differentiated microfiber-hLPs derived from *SFTPC^GFP^* hiPSCs, *SLC34A2^KO^* hiPSCs, PAM#1-4-hiPSCs, and cPAM#1-hiPSCs three days after passaging onto cell culture inserts. (n = 3–4/group). Data are expressed as mean ± SEM; unpaired, two-tailed t-test: *P < 0.05, **P < 0.01. (M) Colorimetric assay of phosphate concentrations in apical medium of monolayer culture in alveolar-differentiated microfiber-hLPs derived from *SFTPC^GFP^*hiPSCs and *SLC34A2^KO^* hiPSCs (n = 8/group from eight independent experiments). Data are expressed as mean ± SEM; two-way repeated measures ANOVA with Turkey’s post-hoc analysis, ns: not significant, **P < 0.01 (N) Colorimetric assay of phosphate concentrations in basolateral medium of monolayer culture in alveolar-differentiated microfiber-hLPs derived from *SFTPC^GFP^*hiPSCs and *SLC34A2^KO^* hiPSCs (n = 8/group from eight independent experiments). Data are expressed as mean ± SEM; two-way repeated measures ANOVA with Turkey’s post-hoc analysis, ns: not significant, **P < 0.01 (O) Diagram illustrating phosphate metabolism in the alveolar region. (P) Bright-field imaging of monolayer culture after changing to medium containing calcium phosphate particles from the apical side. Scale bars, 1 mm. (Q) Quantification of increased or decreased calcium phosphate particle rate from 24 h to 72 h after medium change (n = 6/group). Data are expressed as mean ± SEM; unpaired, two-tailed t-test, **P < 0.01.

RNA-seq analysis was performed on day 21 hLPs, microfiber-hLPs (day 49), alveolar-differentiated microfiber-hLPs (day 63), and their monolayer-passaged counterparts using hiPSCs. Principal component analysis (PCA) revealed that replicates clustered by sample type and pre-passage samples were segregated from monolayer-passaged samples (fig. S4B). Moreover, monolayer day 21 hLPs exhibited a gene expression profile relatively similar to that of pre-passage day 21 hLPs, whereas monolayer alveolar-differentiated microfiber-hLPs had distinct gene expression profiles compared to pre-passage day 21 hLPs (fig. S4, B and C). In particular, microfiber-mediated expansion and differentiation increased the expression of AT1 and AT2 cell markers while decreasing progenitor cell markers (fig. S4D). Even after passage to monolayer culture, the alveolar-differentiated microfiber-hLPs strongly expressed AT2 markers, including *SLC34A2, ABCA3* and *SFTPB*, while *SFTPC* was downregulated. Gene Ontology (GO) enrichment further revealed that terms related to phosphate metabolism, such as “phosphorus metabolic process” and “phosphate-containing compound metabolic process,” were enriched in monolayer alveolar-differentiated microfiber-hLPs compared to monolayer day 21 hLPs.

The proportion of NaPi2b^+^ cells among the total hiPSC-derived cells on cell culture inserts increased after microfiber-mediated expansion and differentiation, indicating their maturation into alveolar lineage (Fig. 3D and fig. S4G). Correspondingly, transmission electron microscopy (TEM) revealed the presence of intercellular adhesion molecules, including tight junctions, adherens junctions, and desmosomes, in the monolayers (fig. S5H). Moreover, microvilli and lamellar bodies, unique to AT2 cells, were observed on the apical side of the monolayer. These results suggested that alveolar-differentiated microfiber-hLPs in monolayer culture had well-established intercellular junctions and maintained apico–basal polarity.

Finally, the monolayer system was employed to analyze phosphate metabolism. A tracer experiment using the radioisotope ^32^P showed higher phosphate uptake from the medium on the apical side into alveolar-differentiated microfiber-hLPs than from the lower chamber (Fig. 3E). Additionally, the phosphate concentration in the apical medium decreased over time, regardless of the initial apical phosphate concentration, while that in the basolateral medium increased (Fig. 3, F and G). In contrast, when using day 21 hLPs or microfiber-hLPs, the phosphate concentration did not change significantly in the apical or basolateral medium (Fig. 3, H and I). Hence, alveolar-differentiated microfiber-hLPs transported phosphate ions from the apical side to the basolateral side.

### Monolayer culture enables disease modeling of pulmonary alveolar microlithiasis

*SLC34A2*, which encodes NaPi2b, is linked to pulmonary alveolar microlithiasis (PAM), a rare autosomal recessive genetic disease characterized by intra-alveolar microliths primarily composed of calcium phosphate (*37*, *38*). To model PAM using hiPSCs, four patients with PAM were recruited (Patients #1–4) (Table. 1). The radiographic images for Patient #1 showed diffuse hyperdense micronodular airspace opacities predominating in the bilateral lower lobes (fig. S5A). The *SLC34A2* variant was determined for each patient, as the variants for Patients #1 and #2 have not been reported (*39*). Disease-specific hiPSCs (PAM#1, #2, #3, and #4-hiPSCs) were generated from each patient’s peripheral blood mononuclear cells (PBMCs; fig. S5B). All PAM-hiPSCs expressed undifferentiated markers and pluripotency (fig. S5C). Subsequently, a gene-corrected hiPSC line (cPAM#1-hiPSCs) was generated from the PAM#1-hiPSCs, as well as a genetic knockout hiPSC line of *SLC34A2* (*SLC34A2^KO^*hiPSCs) from healthy donor-derived *SFTPC^GFP^* hiPSCs using CRISPR/Cas9-mediated gene editing (fig. S5, D to G). Sequencing confirmed the correction of exon 6 mutation in cPAM#1-hiPSCs and a new homozygous frameshift mutation in exon 1 (c.39delC) in *SLC34A2^KO^* hiPSCs. Additionally, PAM#1, #2, #3, and #4-hiPSCs, cPAM#1-hiPSCs and *SLC34A2^KO^* hiPSCs had normal karyotypes (fig. S5H). The isogenic pairs of hiPSC lines were stepwise differentiated and expanded into alveolar-differentiated microfiber-hLPs (Fig. 1F). The proportion of SFTPC-GFP^+^ cells in alveolar-differentiated microfiber-hLPs derived from *SLC34A2^KO^*hiPSCs did not differ from those derived from the parental *SFTPC^GFP^* hiPSCs (fig. S5, I to K). This suggested that *SLC34A2* expression did not affect alveolar lineage differentiation.

**Table 1.**
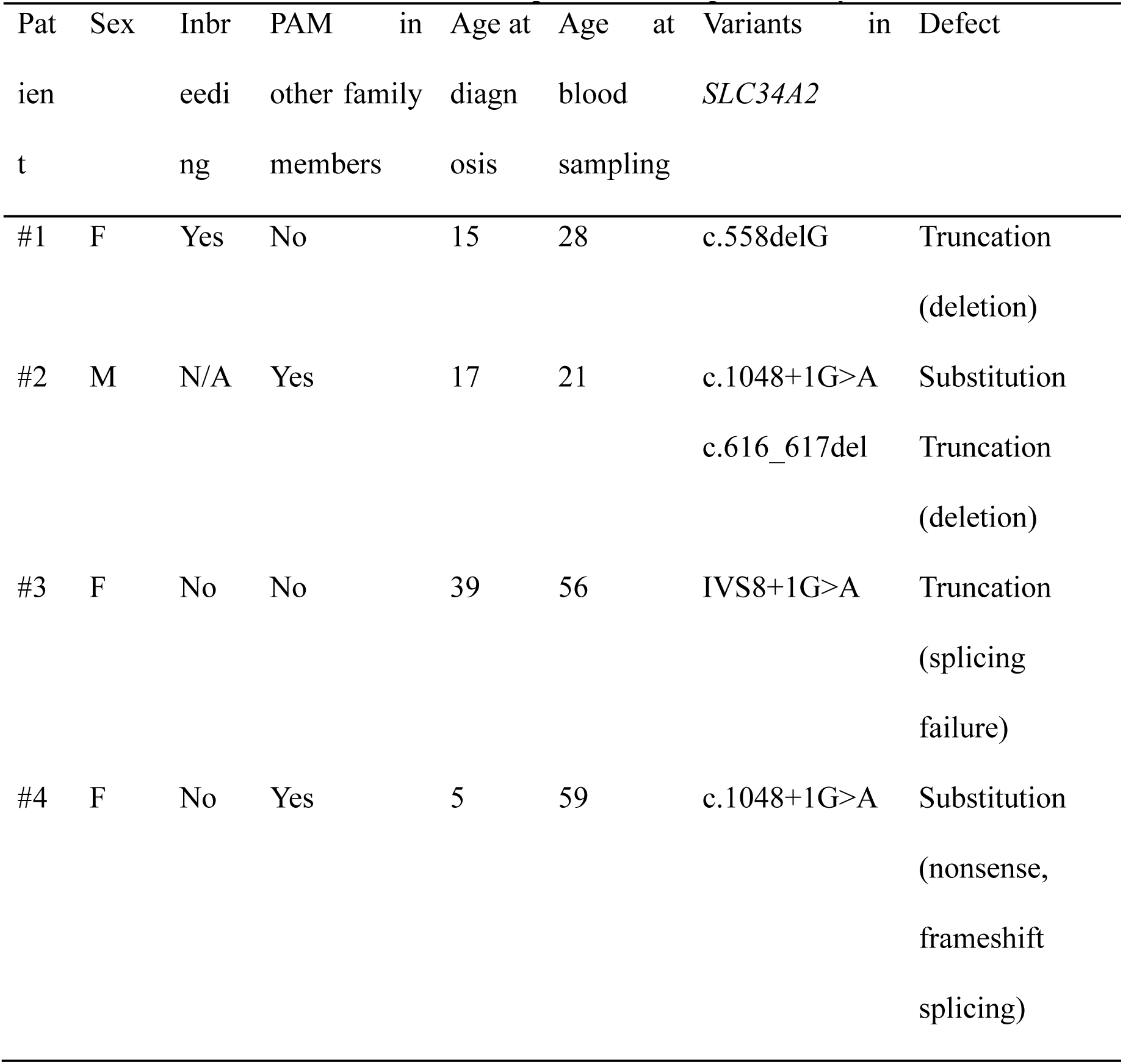
Characteristics of the recruited patients with pulmonary alveolar microlithiasis.

To evaluate NaPi2b’s role in monolayer culture phosphate metabolism, alveolar-differentiated microfiber-hLPs derived from *SFTPC^GFP^*hiPSCs, *SLC34A2^KO^* hiPSCs, PAM#1, #2, #3, and #4-hiPSCs, and cPAM#1-hiPSCs were seeded onto cell culture inserts. Alveolar-differentiated microfiber-hLPs derived from *SFTPC^GFP^* hiPSCs and cPAM#1-hiPSCs expressed NaPi2b on the apical side of the monolayer, whereas those derived from *SLC34A2^KO^* hiPSCs and PAM#1, #2, #3, and #4-hiPSCs did not express NaPi2b (Fig. 3, J and K, and fig. S5L). Tracer experiments demonstrated reduced phosphate uptake by alveolar-differentiated microfiber-hLPs derived from PAM#1, #2, #3, and #4-hiPSCs or *SLC34A2^KO^* hiPSCs (Fig. 3L). Phosphate concentrations were higher in the apical medium and lower in the basaolateral medium of the monolayer derived from *SLC34A2^KO^* hiPSCs than those derived from *SFTPC^GFP^*hiPSCs (Fig. 3, M and N). Moreover, phosphate concentrations in the medium on the apical side of the monolayer derived from cPAM#1-hiPSCs decreased over time, while concentrations did not change significantly on the apical or basolateral sides of monolayers derived from PAM#1, #2, #3, or #4-hiPSCs (fig. S5, M to P). Together, these findings highlight the role of NaPi2b in transporting phosphate from the alveolar space to the basal extracellular space (Fig. 3O).

To recapitulate the PAM environment in the alveolar region, the apical medium was replaced with one containing calcium phosphate particles three days after passaging. Using alveolar-differentiated microfiber-hLPs derived from healthy donor-derived *SFTPC^GFP^* hiPSCs, calcium phosphate particles decreased over time; similar changes were not observed for those derived from *SLC34A2^KO^* hiPSCs (Fig. 3, P and Q). Thus, a novel *in vitro* PAM model was established using alveolar-differentiated microfiber-hLPs.

### Transplanting disease-specific hiPSC-derived hLPs recapitulates the PAM phenotype *in vivo*

To assess the role of AT2 cells in phosphate homeostasis within the alveolar space after transplantation, the expression of genes involved in phosphate metabolism was evaluated using scRNA-seq (Fig. 4, A and B). *SLC34A2* (NaPi2b) was present in all clusters, while *SLC34A1* (NaPi2a) and *SLC34A3* (NaPi2c), which are typically expressed in the kidney (*40*, *41*), were absent in all clusters. *SLC20A1* (PiT-1) and *SLC20A2* (PiT-2), ubiquitously expressed in various tissues (*42*), were also detected in all clusters. *XPR1*, the only known phosphate exporter (*43*), was expressed in all clusters.

**Fig. 4.**
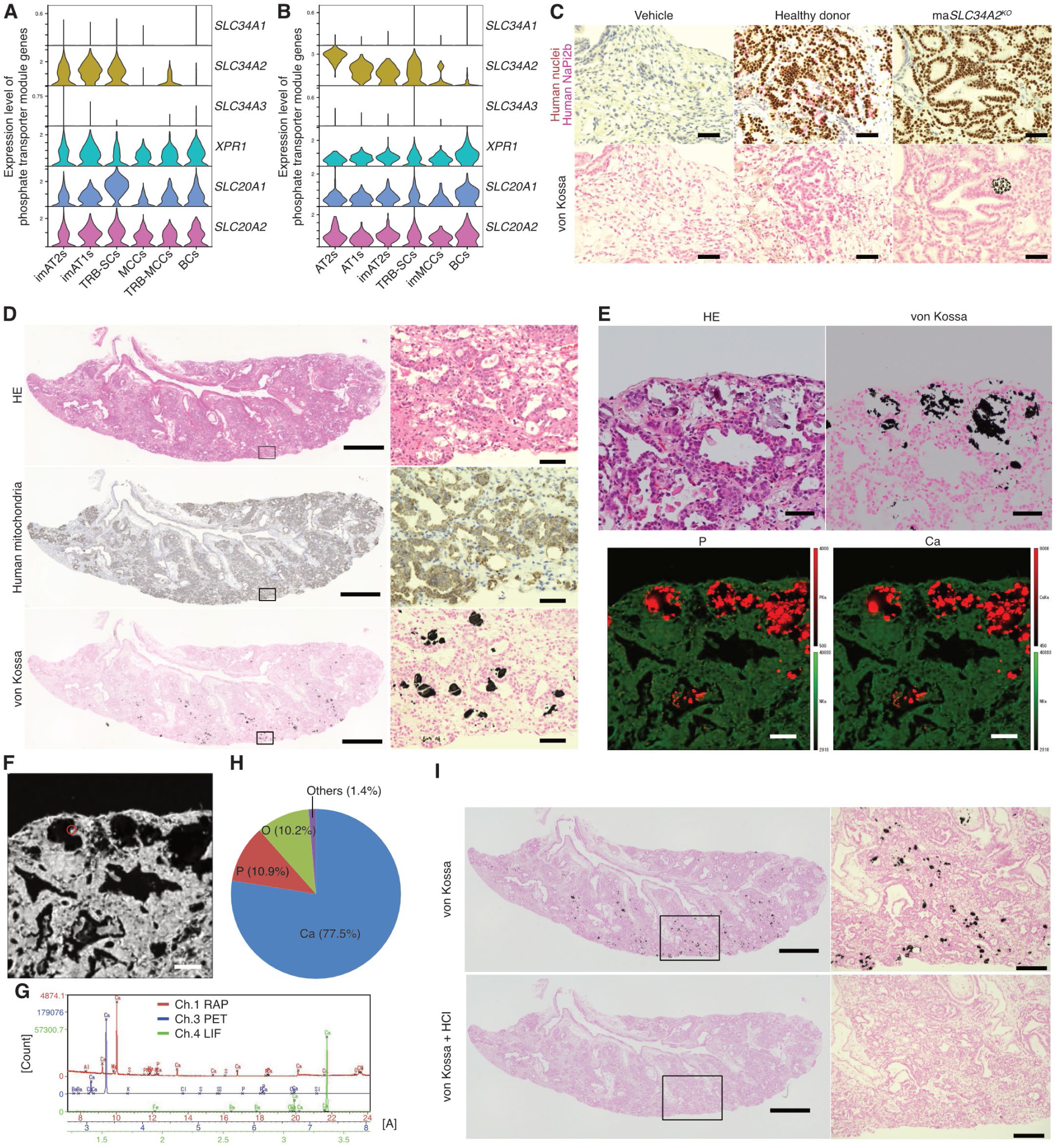
Transplanting disease-specific hiPSC-derived hLPs recapitulates the PAM phenotype *in vivo*. (A, B) Violin plots of phosphate transporter module gene expression (*SLC34A1*, *SLC34A2*, *SLC34A3*, *XPR1*, *SLC20A1,* and *SLC20A2*) across cell clusters identified by scRNA-seq of engrafted day 21 hLPs (A) and engrafted alveolar-differentiated microfiber-hLPs (B) 24 weeks post-transplantation. (C) IHC and von Kossa staining of the serial left lung sections of *NOG-LysM-DTR* mice 24 weeks post-transplantation of day 21 hLPs derived from *mCherry-Akaluc SFTPC^GFP^* hiPSCs and *mCherry-Akaluc SLC34A2^KO^* (ma*SLC34A2^KO^*) hiPSCs. Scale bar, 20 μm. (D) Hematoxylin and eosin (H&E), immunohistochemical (IHC), and von Kossa staining of serial left lung sections from *NOG-LysM-DTR* mice 24 weeks post-transplantation of alveolar-differentiated microfiber-hLPs derived from *mCherry-Akaluc* PAM#1 (maPAM#1)-hiPSCs. Scale bar, 1 mm (Left), 40 μm (Right). (E) Representative light microscopic image and EPMA–WDS image of serial left lung sections from *NOG-LysM-DTR* mice 24 weeks post-transplantation of alveolar-differentiated microfiber-hLPs derived from maPAM#1-hiPSCs. Scale bar, 40 μm. (F) Gray scale image; Scale bar, 40 μm; red circle: region that was one-dimensionally analyzed, with results shown in (G, H). (G) One-dimensional analysis yields element peaks on the curves as detected with WDS crystals: rubidium acid phthalate (RAP, red), pentaerythritol (PET, blue), and lithium fluoride (LiF, green). (H) Elemental composition of the region of interest. Data are expressed as mol (%). (I) Von Kossa staining with or without hydrochloric acid treatment of left lung sections of *NOG-LysM-DTR* mice 24 weeks after transplantation of alveolar-differentiated microfiber-hLPs derived from maPAM#1-hiPSCs. Scale bar, 1 mm (left), 200 μm (right).

To evaluate the utility of lung-humanized mice as a platform for *in vivo* disease modeling, *mCherry-Akaluc* reporter *SLC34A2^KO^*hiPSCs and PAM#1-hiPSC lines were constructed (ma*SLC34A2^KO^* hiPSCs and maPAM#1-hiPSCs, respectively). ma*SLC34A2^KO^* hiPSCs and maPAM#1-hiPSCs had normal karyotypes (fig. S6A). These hiPSC-derived day 21 hLPs or alveolar-differentiated microfiber-hLPs were transplanted into the left lungs of *NOG-LysM-DTR* mice. Von Kossa staining of the left lobe 24 weeks post-transplantation revealed calcium deposits in areas with engrafted cells derived from ma*SLC34A2^KO^*hiPSCs, not *mCherry-Akaluc SFTPC^GFP^* hiPSCs or in vehicle-injected lungs (Fig. 4C). Calcium deposits were detected 24 weeks after transplantation of day 21 hLPs derived from maPAM#1-hiPSCs, but not at earlier time points (fig. S6B). Live imaging and histological analysis of the left lobe confirmed extensive engraftment and calcium deposits 24 weeks after transplantation of alveolar-differentiated microfiber-hLPs derived from maPAM#1-hiPSCs (Fig. 4D and fig. S6C). Elemental analysis of the histological sample with an electron probe microanalyzer-wavelength dispersive spectrometer (EPMA-WDS) revealed deposits primarily composed of calcium, oxygen, and phosphorus (Fig. 4, E to H, and fig. S6, D to G), and to a lesser degree, iron, magnesium, potassium, aluminum, and silica, similar to microliths in patients with PAM (*44*, *45*). This was further confirmed by the dissolution of deposits after treatment with hydrochloric acid (Fig. 4I). Hence, transplanting disease-specific hiPSC-derived hLPs successfully recapitulated the PAM phenotype *in vivo*.

## Discussion

In this study, a human-lung chimeric mouse model was established with human alveolar epithelial cells architecturally and functionally reconstituted and maintained long-term *in vivo*. This EATAS transplantation method minimizes off-target toxin exposure, selectively ablating endogenous AT2 cells and lung-resident macrophages exclusively in the left lobe. This preserves the contralateral lung, avoiding lethal respiratory compromise, and allowing extended observation.

*NOG* mice retain certain immune functions associated with macrophages, mast cells, and neutrophils, causing a low but significant transplanted cell rejection (*46*, *47*). Meanwhile, simultaneously ablating AT2 cells and resident macrophages reduced barriers to transplanted cell engraftment, achieving an ∼82% replacement rate, the highest reported for orthotopic xenotransplantation into the alveolar region.

The results indicated that transplanted alveolar-differentiated microfiber-hLPs generated AT1 and AT2 cells similar to human primary alveolar epithelial cells in transcriptomics and functions. AT2 cells function as stem-like epithelial cells in the alveoli, differentiating into AT1 cells after injury (*25*, *48*–*50*). At 24 weeks of transplantation, AT1 cells with large, flat and thin morphology were observed near AT2 cells, suggesting that the engrafted AT2 cells differentiated further into AT1 cells for regeneration, thereby reconstructing a pseudo-human alveolar unit.

MHC-Ⅱ, which was not previously reported as being expressed in hiPSC *in vitro* culture system, was detected in the engrafted alveolar-differentiated microfiber-hLPs. While AT2 cells contribute to adaptive immune responses by constitutively expressing MHC-Ⅱ, the regulatory signals for MHC-Ⅱ expression in AT2 cells remain unidentified (*30*). A recent study using fetal-lung derived AT2 organoids suggested that immune function cannot be acquired in a cell-autonomous manner *in vitro* (*51*). Indeed, pre-transplant alveolar-differentiated microfiber-hLPs did not express MHC-Ⅱ, and the host microenvironment may have mediated MHC-II expression in engrafted AT2 cells. These findings, along with the secretion of lung-specific proteins by engrafted cells, elevate this platform beyond a structural replacement, allowing for the investigation of alveolar epithelial physiology and metabolism *in vivo*.

Disease modeling validated the system’s versatility. Transplanting disease-specific hiPSCs into the airway reproduces the primary ciliary dyskinesia phenotype (*52*), but no reports have shown the reproducibility of disease phenotypes in the alveolar region. In the current study, the PAM phenotype was reconstructed *in vitro* and *in vivo*. More specifically, *in vitro*, while calcium phosphate particles were retained within the monolayer apical medium of alveolar-differentiated microfiber-hLPs derived from disease-specific-hiPSCs, microliths did not form spontaneously in this medium. In contrast, at least 24 weeks after transplantation of disease-specific hiPSC-derived cells, microliths formed spontaneously in the alveolar space *in vivo*. This difference may be due to the long period required for microlith formation, which cannot be successfully modelled *in vitro,* given the need to change culture medium regularly to retain cell viability. This highlights the limitations of a simplified *in vitro* model for a complex, progressive disease and the importance of *in vivo* models to better capture the full spectrum of disease pathology.

This study has several limitations that warrant discussion. Although macrophage depletion facilitated engraftment, it removed key regulators of tissue repair after injury (*53*–*56*). Recruited monocytes and tissue-resident alveolar macrophages also contribute to microlith clearance (*57*). Future iterations of the model incorporating human immune cells will allow more detailed analysis of epithelial–immune cross-talk in lung regeneration and disease modeling. Additionally, our model only reproduces the epithelial compartments; species-specific differences in endothelium, mesenchyme, and ventilation may influence long-term maturation of transplanted cells. Finally, although the left-lobe-specific niche simplifies imaging, it imposes regional differences in intrathoracic pressure and perfusion that could affect epithelial behavior. Addressing these issues with inducible genetic tools, tissue-engineered scaffolds, and larger-animal hosts will broaden the translational relevance of the platform.

In conclusion, we established a novel model of hiPSC-derived alveolar epithelial cell transplantation that achieves durable and efficient engraftment. This chimeric lung platform precisely captures the architectural, functional, and metabolic complexity of human alveolar epithelium *in vivo* and recapitulates the PAM phenotype. Beyond providing a proof-of-concept for regenerative therapy, this system serves as a bridge between existing experimental models and first-in-human studies, facilitating the development of new therapeutics for intractable lung diseases.

## Acknowledgments

We are grateful to the patients and their family. We thank all of the members at the Gotoh lab (CiRA, Kyoto University) for supporting general experiments; N. Suzuki (Central Institute for Experimental Medicine and Life Science) for supporting animal experiments; M. Goto (Central Institute for Experimental Medicine and Life Science) for embryo transfer and production of *NOG-LysM-DTR* mice; M. Mochizuki (Central Institute for Experimental Medicine and Life Science) for histopathological assays. K. Kohno and M. Tanaka (RIKEN) for *LysM-DTR* mice. A. Hotta (CiRA, Kyoto University) for CRISPR-related plasmids; K. Okita, K. Deguchi, and S. Sakurai (CiRA, Kyoto University) for bulk RNA-seq and single-cell RNA-seq analysis; Single-Cell Genome Information Analysis core (SignAC) in ASHBi for the RNA sequence analysis; K. Okamoto-Furuta and T. Katsuno (Division of Electron Microscopic Study, Center for Anatomical Studies, Kyoto University) for supporting electron microscopy; M. Horie (Radioisotope Research Center, Kyoto University) for supporting RI experiments; K. Takakura (Live Imaging Center, Kyoto University) for supporting lightsheet microscopy imaging; M. Narita (CiRA, Kyoto University) for proteomic analysis; G. Ritter (Ludwig Institute for Cancer Research, New York City, USA) for providing anti-NaPi2b antibody (MX35); K Woltjen (CiRA, Kyoto University) for providing plasmid vectors (pXAT2 and pAAVS1-Nst-MCS). The pcDNA3 Venus-Akaluc was provided by the RIKEN BRC through the National BioResource Project of the MEXT, Japan (cat. RDB15781). A part of the fluorescence studies was performed at Medical Research Support Center, Kyoto University. A part of *in vivo* imaging studies was conducted through the CORE Program of the Radiation Biology Center, Kyoto University. A part of the histopathological assays was conducted in the Center for Anatomical, Pathological and Forensic Medical Research, Kyoto University. Illustrations were created by BioRender.

## Supplementary Materials

**fig. S1.**
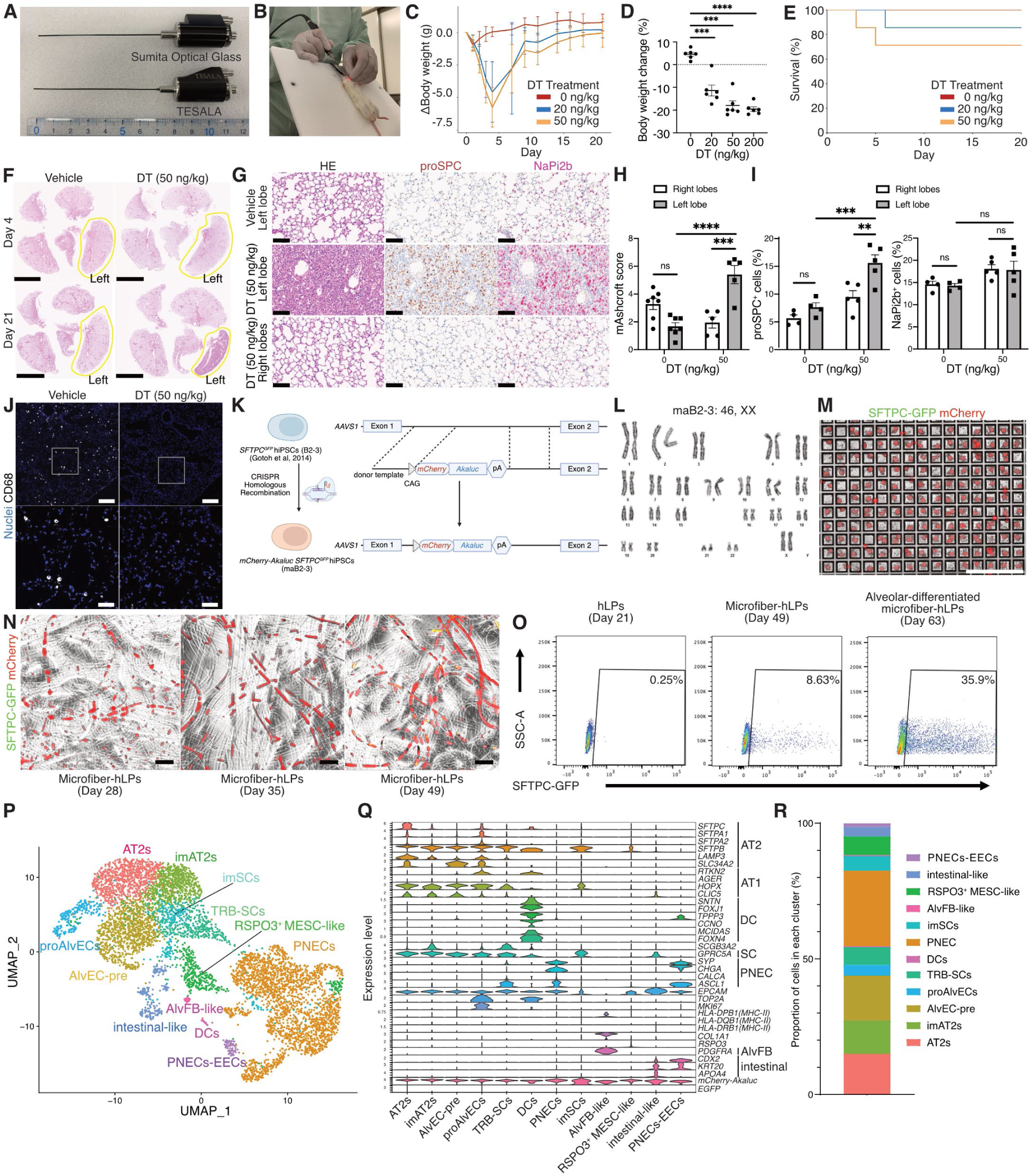
Validation of *NOG-LysM-DTR* mice and differentiation of luminescent-fluorescent reporter hiPSCs into alveolar-differentiated microfiber-hLPs for transplantation. (A) Endoscopes used in this study. (B) Performance of the endoscope-assisted transtracheal administration system (EATAS). (C) Weight loss associated with DT treatment (mean ± SEM, n = 6 mice/group). (D) Body weight changes four days after DT treatment (mean ± SEM, n = 6 mice/group); one-way analysis of variance (ANOVA) with Turkey’s post-hoc analysis, ***P < 0.001, ****P < 0.0001. (E) Survival curve associated with DT treatment (n = 6 mice/group). (F) Hematoxylin and eosin (H&E) staining of lung sections 4 or 21 days after saline or DT treatment. Scale bar, 5 mm. (G) H&E staining and immunohistochemical (IHC) analysis of lung sections three weeks after vehicle or DT treatment. Scale bar, 100 μm. (H) Assessment of modified Ashcroft score three weeks after DT treatment (mean ± SEM, n = 5–7/group); two-way repeated measures ANOVA with Sidak’s post-hoc analysis. ns: not significant, ***P < 0.001, ****P < 0.0001. (I) Assessment of proSPC^+^ cells and NaPi2b^+^ cells three weeks after DT treatment (mean ± SEM, n = 4–5/group); two-way repeated measures ANOVA with Sidak’s post-hoc analysis; ns: not significant, **P < 0.01, ***P < 0.001. (J) Immunofluorescence staining of lung sections four days after vehicle or DT treatment. Scale bars, 200 μm and 50 μm. (K) Schematic of the strategy used to construct the *mCherry-Akaluc SFTPC^GFP^*hiPSC line (maB2-3) by genome editing. (L) Karyotype of the maB2-3 hiPSC line. (M) Live image of hLPs 2 days after seeding on Elplasia plate. Scale bar, 1 mm. (N) Live imaging of microfiber-hLPs on days 28, 35, and 49. Scale bar, 1 mm. (O) Representative flow cytometric analysis of day 21 hLPs, microfiber-hLPs (day 49), and alveolar-differentiated microfiber-hLPs (day 63). (P) UMAP plot of scRNA-seq transcriptomes of pre-transplant alveolar-differentiated microfiber-hLPs. Abbreviations used in the figure include imAT2s (immature alveolar type 2 cells), AlvEC-pre (alveolar epithelial cell precursor cells), proAlvECs (proliferative alveolar epithelial cells), TRB-SCs (terminal and respiratory bronchiole-secretory cells), DCs (deuterosomal cells), PNEC (pulmonary neuroendocrine cell), imSCs (immature secretory cells), AlvFB (alveolar fibroblast), MESC (mesenchymal cell), and EECs (enteroendocrine cells). (Q) Violin plot of selected marker gene expression across clusters. (R) Bar plot showing the proportion of cells in alveolar-differentiated microfiber-hLPs.

**fig. S2.**
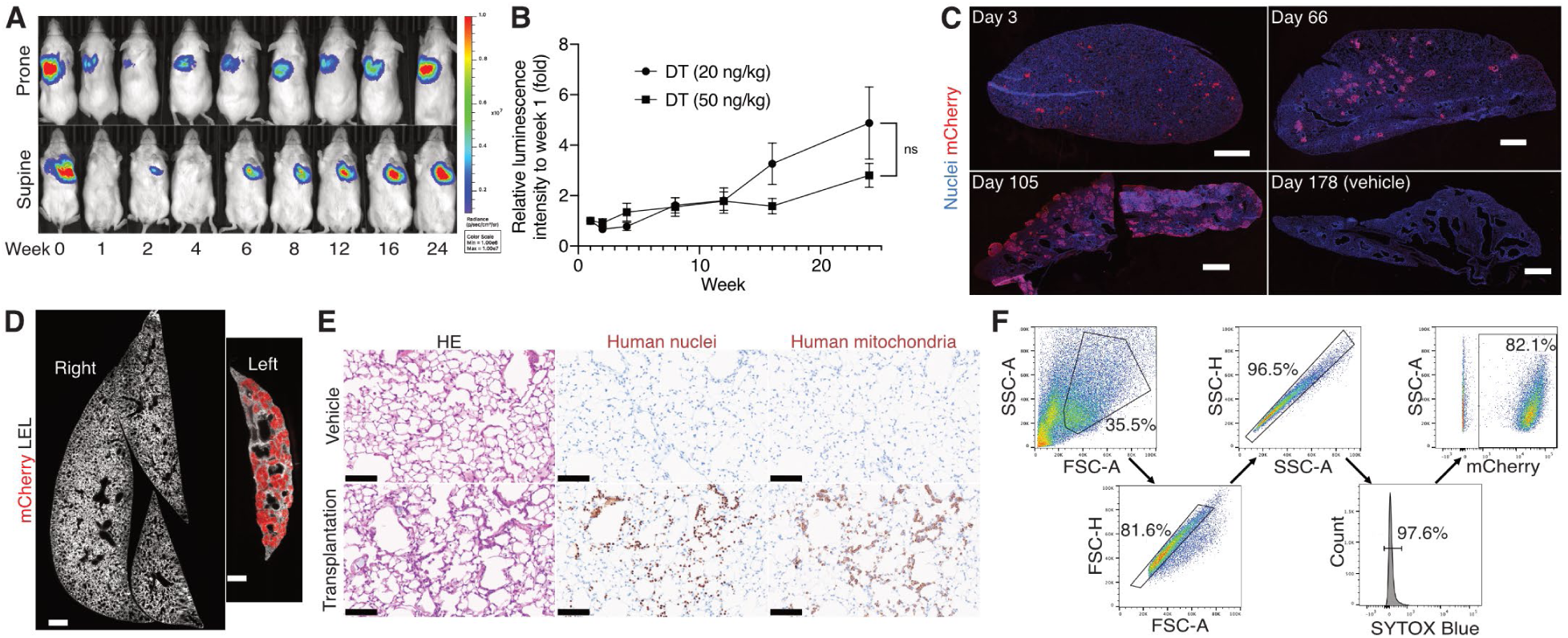
Long-term engraftment of hLPs in the left lobe of *NOG-LysM-DTR* mice. (A) Representative images of *in vivo* luminescence imaging of a mouse transplanted with alveolar-differentiated microfiber-hLPs following injury with diphtheria toxin (DT) 50 ng/kg. (B) Relative bioluminescence intensity for 24 weeks post-transplantation of alveolar-differentiated microfiber-hLPs following injury with 20 ng/kg (n = 7) or 50 ng/kg (n = 8) DT, with a luminescence intensity of 1 at week one (mean ± SEM); two-way repeated measures analysis of variance (ANOVA) with Sidak’s post-hoc analysis; ns: not significant. (C) Immunofluorescence imaging of the left lobe of *NOG-LysM-DTR* mice after alveolar-differentiated microfiber-hLP transplantation; Scale bars, 1 mm. (D) Representative cross sections of cleared mouse left and right lobes 24 weeks post-transplantation of day 21 hLPs. Scale bars, 1 mm. (E) Hematoxylin and eosin (H&E) staining and immunohistochemical (IHC) analysis of lung sections 24 weeks post-transplantation of alveolar-differentiated microfiber-hLPs derived from *SLC34A2^KO^* hiPSCs. Scale bar, 1 mm. (F) FACS gating strategy for isolating mCherry^+^ human cells in the single-cell suspensions of *NOG-LysM-DTR* mouse lungs.

**fig. S3.**
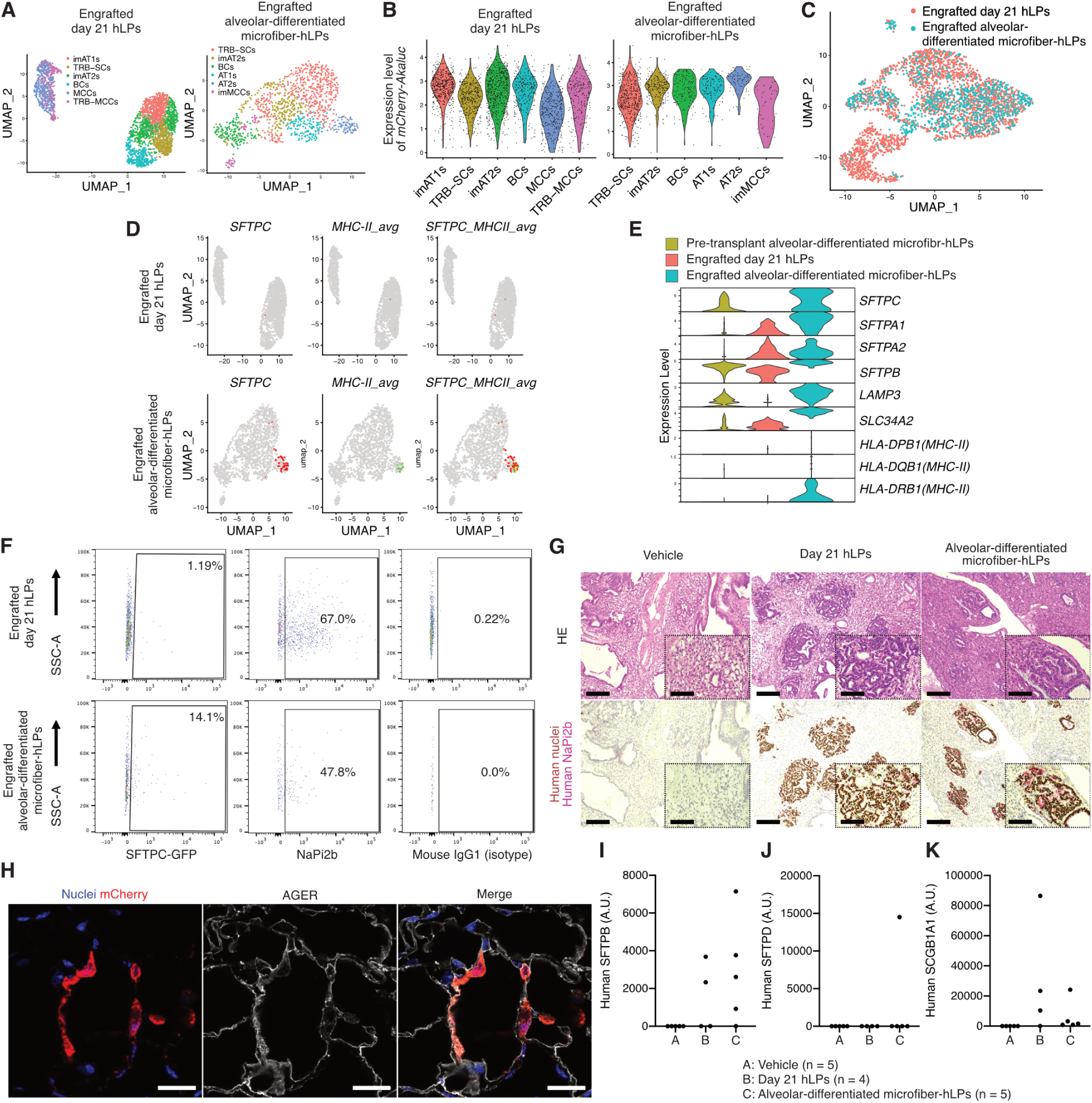
Transplanted hLPs differentiate into alveolar epithelial cells *in vivo*. (A) UMAP plot of scRNA-seq transcriptomes of the engrafted hLPs in each condition. Abbreviations used in the figure include imAT2s (immature alveolar type 2 cells), imAT1s (immature alveolar type 1 cells), and imMCCs (immature multiciliated cells). (B) Violin plots showing Z scores of *mCherry-Akaluc* in engrafted day 21 hLPs and alveolar-differentiated microfiber-hLPs. (C) Integrated UMAP plot of scRNA-seq transcriptomes of the engrafted day 21 hLPs and alveolar-differentiated microfiber-hLPs. (D) Feature UMAP of *MHCⅡ* and *SFTPC* expression. (E) Stacked violin plots of AT2 cell marker gene expression across AT2 clusters, split by condition: pre-transplant alveolar-differentiated microfiber-hLPs (left), engrafted day 21 hLPs (middle), and engrafted alveolar-differentiated microfiber-hLPs (right) (F) Representative flow cytometric analysis of SFTPC-GFP^+^ cells and NaPi2b^+^ cells for mCherry^+^ cells. (G) Hematoxylin and eosin (H&E) staining and immunohistochemical (IHC) analysis of *NOG-LysM-DTR* mice left lung sections 24 weeks post-transplantation of day 21 hLPs. Scale bars, 200 μm and 100 μm. (H) Confocal microscopic images depicting mCherry^+^ engrafted lesion in lungs of *NOG-LysM-DTR* mice 24 weeks post-transplantation of alveolar-differentiated microfiber-hLPs. Scale bar, 20 μm. (I–K) Abundance of human SFTPB (I), SFTPD (J), and SCGB1A1 (K) across BALF samples from mice 24 weeks post-transplantation of vehicle (n = 5), day 21 hLPs (n = 4), and alveolar-differentiated microfiber-hLPs (n = 5). Values are presented in protein area (A.U.), derived from mass spectrometry-based proteomics.

**fig. S4.**
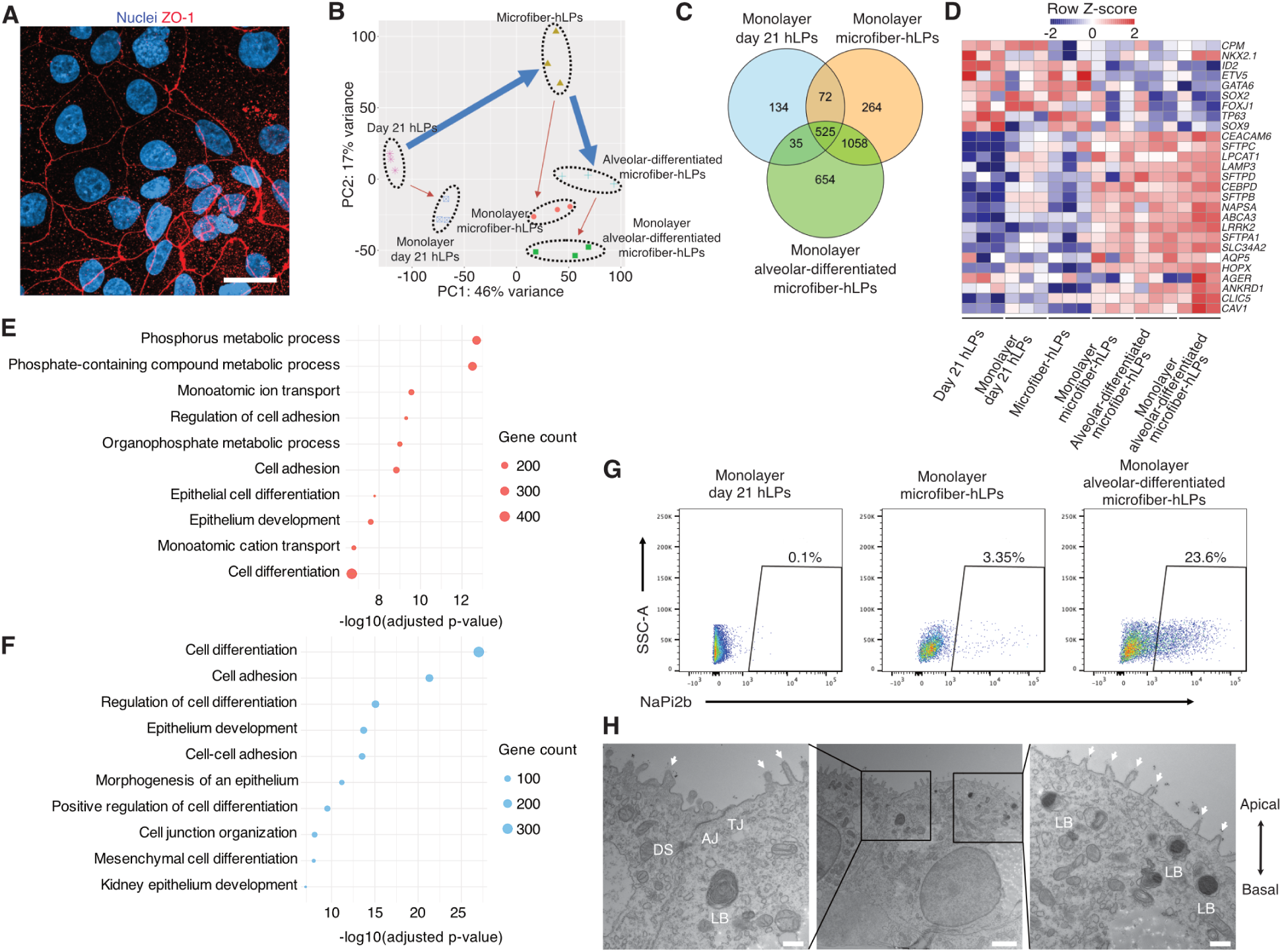
Monolayer alveolar-differentiated microfiber-hLP cultures recapitulate barrier integrity and cell polarity. (A) Immunofluorescence image of reseeded alveolar-differentiated microfiber-hLPs derived from *SFTPC^GFP^* hiPSCs grown on a cell culture insert for three days. Scale bar, 20 µm. (B) Principal component analysis (PCA) of six samples (n = 3/condition) showing global transcriptomic variance (%) of PC1 and PC2. Each plot indicates the transcriptomes of day 21 hLPs, three days after reseeding on cell culture inserts; microfiber-hLPs, three days after reseeding on cell culture inserts; alveolar-differentiated microfiber-hLPs, three days after reseeding on cell culture inserts (n = 3/condition). (C) Venn diagram of differentially expressed genes (DEGs) upregulated in monolayers of reseeded day 21 hLPs, microfiber-hLPs, and alveolar-differentiated microfiber-hLPs compared with before reseeding of day 21 hLPs. The threshold for upregulation was log2 (fold-change) > 1, adjusted P value < 0.05. (D) Heatmap of row-normalized expression of key lung progenitor markers, AT2 markers, and AT1 markers. (E) Representative Gene Ontology (GO) biological processes in upregulated DEGs (log2 fold change [FC] > 1, FDR < 0.05) in monolayered alveolar-differentiated microfiber-hLPs compared with monolayered day 21 hLPs. (F) Representative GO biological processes in downregulated DEGs (log2FC < −1, FDR < 0.05) in monolayered alveolar-differentiated microfiber-hLPs compared with monolayered day 21 hLPs. (G) Representative flow cytometric analysis of day 21 hLPs, microfiber-hLPs, and alveolar-differentiated microfiber-hLPs derived from *SFTPC^GFP^* hiPSCs three days after reseeding onto cell culture inserts. (H) Transmission electron micrograph of alveolar-differentiated microfiber-hLPs derived from *SFTPC^GFP^* hiPSCs three days after reseeding onto cell culture inserts. TJ, tight junction; AJ, adherence junction; DS, desmosome; LB, lamellar bodies. Arrowhead, microvilli. Scale bars, 500 nm (left & right) and 2 µm (middle).

**fig. S5.**
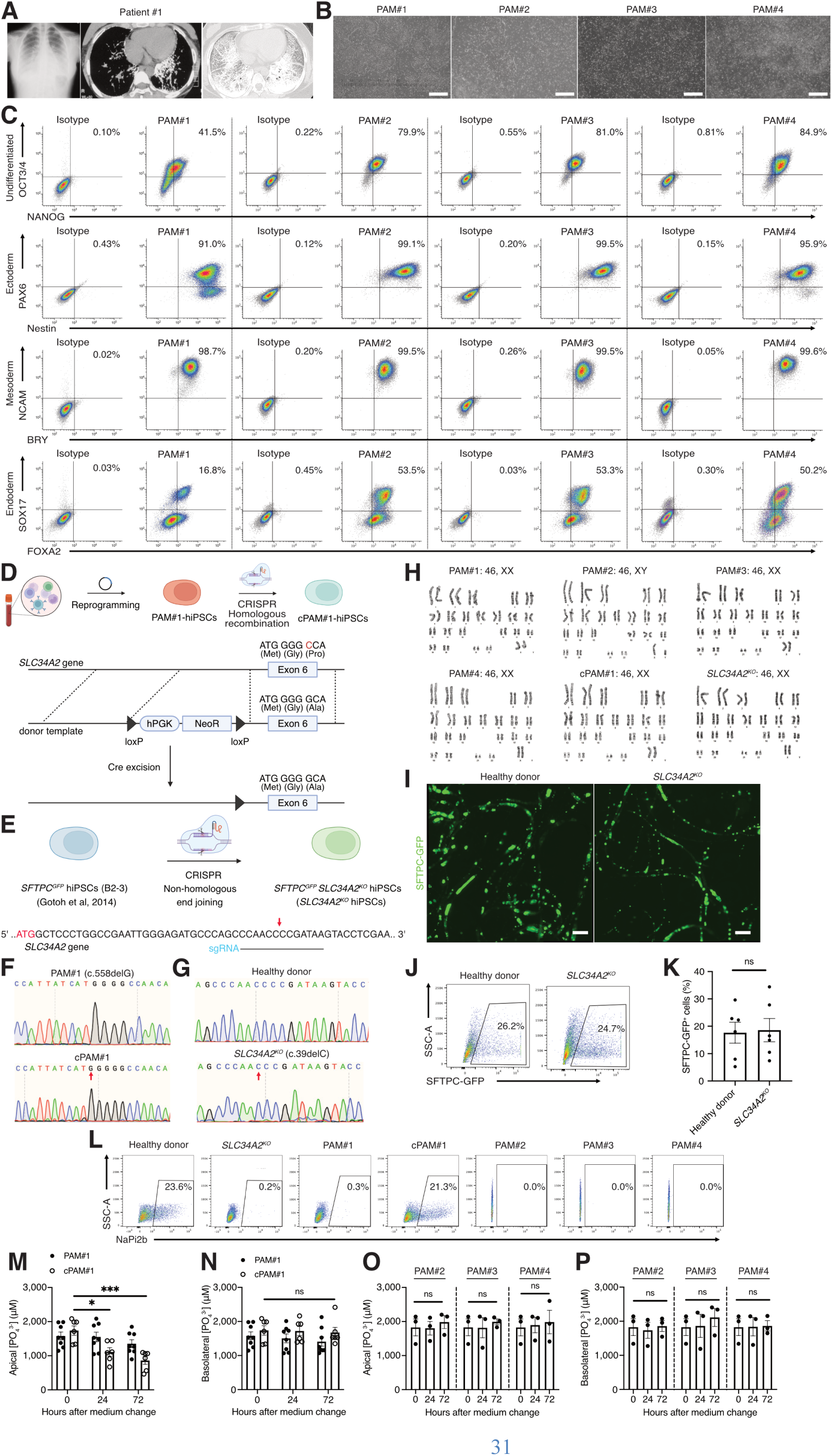
Generation and validation of disease-specific PAM hiPSCs. (A) Representative chest images of patient #1 with PAM. (B) Bright field images of undifferentiated hiPSC lines derived from Patient #1–4. Scale bar, 500 μm. (C) Flow cytometric analysis of undifferentiated cell markers for each hiPSC line and their trilineage-differentiation potency. (D) Schematic of the strategy to generate gene-corrected hiPSCs from PAM#1-hiPSCs (cPAM#1-hiPSCs). (E) Schematic of the strategy to generate *SLC34A2* knockout hiPSCs from *SFTPC^GFP^*hiPSC line (*SLC34A2^KO^* hiPSCs). (F) Sequence data of *SLC34A2* Exon 6 in PAM#1-hiPSCs and cPAM#1-hiPSCs. (G) Sequence data of *SLC34A2* Exon 1 in *SFTPC^GFP^* hiPSCs and *SLC34A2^KO^* hiPSCs. (H) Karyotypes of PAM-hiPSCs (PAM#1–4), cPAM#1-hiPSCs and *SLC34A2^KO^* hiPSCs. (I) Live imaging of alveolar-differentiated microfiber-hLPs derived from *SFTPC^GFP^* hiPSCs and *SLC34A2^KO^* hiPSCs. Scale bar, 1 mm. (J) Representative flow cytometric analysis of alveolar-differentiated microfiber-hLPs (day 63) derived from *SFTPC^GFP^* hiPSCs and *SLC34A2^KO^* hiPSCs, respectively. (K) Flow cytometric quantification of the proportion of PC-GFP^+^ cells in alveolar-differentiated microfiber-hLPs (mean ± SEM, n = 6/group); paired t-test, ns: not significant. (L) Representative flow cytometric analysis of alveolar-differentiated microfiber-hLPs derived from *SFTPC^GFP^* hiPSCs, *SLC34A2^KO^* hiPSCs, PAM#1–4-hiPSCs, cPAM#1-hiPSCs three days after reseeding onto cell culture inserts. (M) Colorimetric assay of phosphate concentrations in the apical medium of monolayer cultures 0 h, 24 h, and 72 h after medium change in alveolar-differentiated microfiber-hLPs derived from PAM#1-hiPSCs and cPAM#1-hiPSCs (n = 6–8/group). Data are expressed as mean ± SEM; one-way repeated measures analysis of variance (ANOVA) with Turkey’s post-hoc analysis, ns: not significant, *P < 0.05, ***P < 0.001. (N) Colorimetric assay of phosphate concentrations in the basolateral medium of monolayer cultures 0 h, 24 h, and 72 h after medium change in alveolar-differentiated microfiber-hLPs derived from PAM#1-hiPSCs and cPAM#1-hiPSCs (n = 6–8/group). Data are expressed as mean ± SEM, one-way repeated measures ANOVA with Turkey’s post-hoc analysis, ns: not significant. (O) Colorimetric assay of phosphate concentrations in the apical medium of monolayer cultures 0 h, 24 h, and 72 h after medium change in alveolar-differentiated microfiber-hLPs derived from PAM#2, #3, #4-hiPSCs (n = 3/group). Data are expressed as mean ± SEM; one-way repeated measures ANOVA with Turkey’s post-hoc analysis, ns: not significant. (P) Colorimetric assay of phosphate concentrations in the basolateral medium of monolayer cultures 0 h, 24 h, and 72 h after medium change in alveolar-differentiated microfiber-hLPs derived from PAM#2, #3, #4-hiPSCs (n = 3/group). Data are expressed as mean ± SEM; one-way repeated measures ANOVA with Turkey’s post-hoc analysis, ns: not significant.

**fig. S6.**
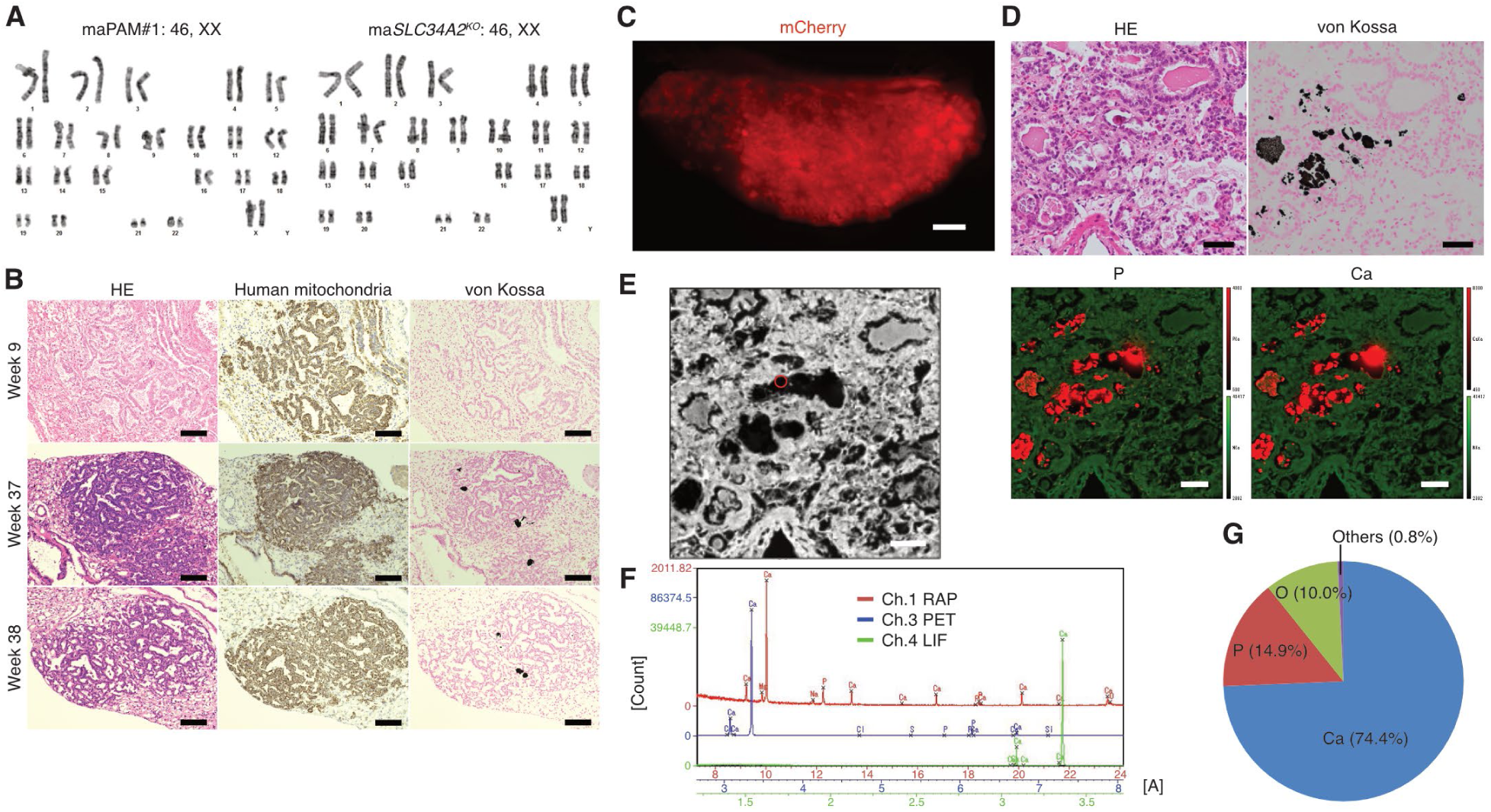
Transplantation of disease-specific hiPSC-derived hLPs recapitulates the phenotype of PAM *in vivo*. (A) Karyotypes of maPAM#1-hiPSCs and ma*SLC34A2^KO^* hiPSCs. (B) Hematoxylin and eosin (H&E) staining, immunohistochemical (IHC) analysis, and von Kossa staining of serial left lung sections from *NOG-LysM-DTR* mice at 9 weeks (upper), 37 weeks (middle), and 38 weeks (lower) post-transplantation of day 21 hLPs derived from maPAM#1-hiPSCs. Scale bar, 100 μm. (C) Representative live imaging of the left lobe of *NOG-LysM-DTR* mice 24 weeks post-transplantation of alveolar-differentiated microfiber-hLPs derived from maPAM#1-hiPSCs. Scale bar, 1 mm. (D) Representative light microscope images and EPMA–WDS images of the serial left lung sections from *NOG-LysM-DTR* mice 24 weeks post-transplantation of alveolar-differentiated microfiber-hLPs derived from maPAM#1-hiPSCs. Scale bar, 40 μm. (E) Gray scale image. Scale bar, 40 μm; red circle: one-dimensionally analyzed region, with results shown in (F, G) (F) One-dimensional analysis yields element peaks on the curves as detected with WDS crystals: rubidium acid phthalate (RAP, red), pentaerythritol (PET, blue), and lithium fluoride (LiF, green). (G) Elemental composition of region of interest. Data are expressed as mol (%).

**Movie S1.**

Part of the endoscope-assisted transtracheal administration system (EATAS) procedure. The outer cylinder of the indwelling needle was placed into the left main bronchus under the endoscope.

**Movie S2.**

3D image of a cleared left lobe 24 weeks after transplanting day 21 hLPs rendered on Imaris. SFTPC-GFP (Green), mCherry (Red), Lycopersicon Esculentum Lectin (LEL; White). Scale bar, 1 mm.

